# Geostatistical mapping of transboundary cattle disease risks in Ethiopia

**DOI:** 10.64898/2026.02.27.708445

**Authors:** Solomon Gizaw, Hiwot Desta, Barbara Wieland, Theodore Knight-Jones

## Abstract

Ethiopia experiences devastating economic losses from an ongoing endemic burden of trans-boundary animal diseases (TADs). TADs are highly transmissible infectious diseases of animals, often able to spread rapidly with significant economic and public health consequences. Contagious bovine pleuropneumonia (CBPP), foot-and-mouth disease (FMD), and lumpy skin disease (LSD) are among the global priority TADs for cattle. In this study, we used responses from a survey about cattle disease delivered to livestock keepers across Ethiopia. We used generalized additive mixed models applying neighborhood cross-validation method which accounts for spatial dependence in the data to investigate the spatially variable relationships between bioclimatic variables and distribution of CBPP, FMD, and LSD in Ethiopia. We also developed model-based risk maps of these diseases using a geostatistical kriging method to guide knowledge-based decision making. The results show the risks of CBPP vary with altitude and relative humidity, risks of FMD with temperature and relative humidity, and of LSD with temperature and precipitation. The gaussian spatial smooth terms are all significant. The maps are produced using rigorous statistical analysis with very low prediction errors and can thus be considered reliable. Our results have implications for the impacts of climate change, and the vulnerability of communities in high-risk areas. The risk maps illustrate how such maps contribute to climate-informed disease early warning systems.

## 1 Introduction

Livestock diseases are a major constraint on the productivity of Ethiopian livestock, and the livelihoods of livestock keeping households and the wider livestock sector, impacting food security. Livestock are particularly important in Ethiopia, accounting for 20% of GDP, and with 18 million pastoralists entirely dependent on livestock. Ethiopia experiences devastating economic losses from an ongoing endemic burden of trans-boundary animal diseases (TADs), with about half of potential livestock production lost to animal disease burden (Jemberu *et al, In Press)*. TADs are highly transmissible infectious disease of animals, often able to spread rapidly with significant economic and public health consequences (Clemmons et al., 2021). Contagious bovine pleuropneumonia (CBPP), foot-and-mouth disease (FMD), lumpy skin disease (LSD), peste des petites ruminants (PPR), contagious caprine pleuropneumonia (CCPP), sheep and goat pox (SGP), and brucellosis are among the global priority diseases for cattle and small ruminants (Seyoum & Teshome, 2017).

Information on the geographic distribution of and environmental variables associated with livestock diseases is important for their prevention, control, and where feasible, eradication. Studies on the spatial distribution of livestock diseases in Ethiopia are very limited. Asrat et al. (2007) used a geographic information system risk modeling approach considering a combination of altitude, temperature, and land slope to estimate the potential spatial distribution of Fasciola species. Similarly, Yilma and Malone (1998) developed a GIS forecast model based on moisture and thermal conditions to assess the risk of *Fasciola gigantica* in Ethiopia. Menghistu et al. (2018) mapped important zoonotic diseases in the Tigray region of Ethiopia. But these three studies did not use model-based geostatistical methods. To progress with disease control, we need to better understand the spatial and temporal distribution of disease risks and the underlying risk factors and spatial process that drive the major livestock diseases in Ethiopia.

In low-resource settings, disease prevalence mapping relies on empirical prevalence data from a finite, often spatially sparse, set of surveys of communities within the region of interest, possibly supplemented by remotely sensed images that can act as proxies for environmental risk factors (Diggle and Giorgi, 2015). A standard geostatistical model for data of this kind is a generalized linear mixed model (GLM) with binomial error distribution, logistic link, a combination of explanatory variables and a Gaussian spatial stochastic process in the linear predictor (Diggle and Giorgi, 2015). For situations where the relationships between response and explanatory variables is non-linear, an alternative efficient method to GLM is the generalized additive model (GAM) and GAMM for mixed models. GAM (Hastie and Tibshirani, 1986, 1990) is a GLM with a linear predictor involving a sum of smooth functions of covariates. The model allows for flexible specification of the dependence of the response on the covariates, specifying the model only in terms of ‘smooth functions’, rather than detailed parametric relationships (Wood, 2017).

Disease risk maps are developed using Ecological Niche Models (ENM, e.g. Li et al., 2023), spatial models, or combined spatial and some elements of ENM, mainly environmental variables. The latter approach enables us to investigate and control for the spatial structure of observations, covariates, and model residuals, associating disease occurrence data and environmental variables to predict a disease’s potential geographic distribution, and understand the environmental conditions that influence a disease’s presence. Such approaches are used for forecasting, such as predicting future distributions under climate change. However, the challenge with this approach is spatial dependence arising from inherent spatial structure in some data. Undesirable outcomes of such underlying dependences include serious underestimation of predictive error, dependence structures in the data persisting with dependence structures in model residuals violating the assumption of independence. This provides ample opportunity for model overfitting with non-causal predictors, and because predictor variables are often correlated with underlying dependence structures (e.g. climate with space), models may use predictors to overfit the residual dependence structure and thereby remove it, partially or completely (Roberts et al., 2017). A recent method developed by Wood (2025), namely the neighborhood cross-validation method, can be used to deal with this problem of unmodelled short range autocorrelation.

There is a great need for risk maps to inform national and regional livestock disease control (REF), however, in many LMICs, including Ethiopia very limited information is available on TADs distribution. Cost-effective disease control must be risk based, with geographic targeting of high-risk groups essential for targeting hot spots in transmission chains and areas of high disease risk and burden. This is particularly true in settings, like Ethiopia, where the resources available for livestock disease control are limited. There is also a great need to improve our understanding of how climate variables affect disease risk, with forecasting of geographic changes in disease distribution with climate change a priority livestock climate adaptation measure, especially in areas already experiencing massive climate variation, such as Ethiopia.

In this study, we modelled the disease experience of livestock keepers across Ethiopia collected in questionnaire surveys. Data generated from questionnaire surveys could be considered suspect with concerns for the accuracy due to respondents’ limited ability to diagnose specific diseases, potential for exaggerating or sometimes hiding disease impact, and potential for recall bias when recalling historic events. Unreliability/inaccuracy of survey data could also arise from binomial sampling error (Diggle and Giorgi, 2015). The conventional data used in disease risk analysis in Ethiopia are from serological tests and outbreak reports. However, there are often massive biases in these data in many countries, especially LMICs, due to under-reporting and inadequate and varying coverage of disease surveillance systems, as well as the unaddressed complexity of interpreting sero-prevalence in surveys of animals of different ages and time at risk. Furthermore, in many LMICs including Ethiopia, participatory methods of disease surveillance have often been found to provide a much more accurate picture of disease distribution and epidemiology. In this study we conducted geostatistical mapping of three of the most important transboundary diseases in Ethiopia (FMD, LSD and CBPP). The aim was to quantify the spatially variable relationships between environmental variables and the disease distribution in Ethiopia and develop model-based risk maps of these diseases to guide knowledge-based decision making.

## 2 Materials and Methods

### 2.1 DATA

The data for this study were obtained from household surveys of five livestock development projects, namely the Drought Resilience of Sustainable Livelihood program DRSLPI in 2012 and DRSLPII in 2014, the Regional Pastoral Livelihoods Resilience Project (RPLRP) in 2015, and the Livestock and Fishery Sector Development Project (LFSDP) in 2018, and the Health of Ethiopian Animals for Rural Development (HEARD) project in 2019. DRSLP, RPLRP and LFSDP projects were implemented by the Ministry of Agriculture of Ethiopia (MoA). The HEARD project was implemented by the MoA with the International Livestock Research Institute (ILRI) as a key partner. All the five surveys were conducted by ILRI. Both purposive and stratified clustered sampling approaches were used to obtain representative samples for the household surveys. Sampling was stratified at different stages, namely by livelihood zones (pastoral, agropastoral and mixed crop-livestock systems) and administrative zones at hierarchical levels of “regions”, “zones”, “*woredas*” (districts) and *Kebeles* (the smallest administrative unit in Ethiopia). Regions were selected purposively as per the projects’ aims and design. *Woredas* within regions and *kebeles* within *woredas* were selected randomly stratified by livelihood zones. Households were selected randomly considering gender, age and livestock holdings. All the five surveys followed similar protocols as described in detail by Gebremedhin et al. (2017). The surveys were conducted by trained enumerators in the presence of local veterinarians who helped translate veterinary disease names into the local languages.

#### Response variable

In this study we considered three important transboundary cattle diseases (CBPP, contagious bovine pleuropneumonia; FMD, foot and mouth disease; and LSD, lumpy skin disease) out of the twelve cattle diseases surveyed in the five surveys (Gizaw et al., 2021a; 2020). Livestock keeping households were asked whether their cattle herds were affected by each of the three diseases in the last 12 months preceding the survey time and the proportion of each household’s herd affected, either 0-15%, 16-50%, 51-75%, 76-99%, or 99-100%. Respondents with empty response entries were considered as reporting that none of their animals were affected. The dataset consisted of 3340 data points for analysis. The percentages of respondents who did not report disease outbreaks, and those who reported 0-15%, 16-50%, 51-75%, 76-99%, and 99-100% of their herds affected were 76.3%, 13.8%,6.1%, 3.1%, 0.5% and 0.2% for CBPP; 80.6%, 12.1%, 4.6%, 1.8%, 0.5% and 0.4% for FMD; and 76.4%, 13.3%, 7.2%, 2.6%, 0.3% and 0.2% for LSD. As our aim was to map the geographic distribution of the diseases and not within-herd prevalence, the analysis was done at the herd level by defining the response variable as a binary response of disease absence or presence (presence class consisting of responses of disease outbreaks). However, we also defined the response as ordinal variable with four classes of None (no outbreak reported), Low (.0-15%), Medium (16-50%), and High (51-100% of the herds affected). We then compared the two response definitions based on model fit criteria. The proportion of ‘No outbreak’ and ‘outbreak’ responses in the two-level response for CBPP, FMD, and LSD datasets were 0.763 and 0.237, 0.806 and 0.194, and 0.764 and 0.236, respectively. The study area and sampling points (households) are shown in Fig. 1.

**Fig. 1.**
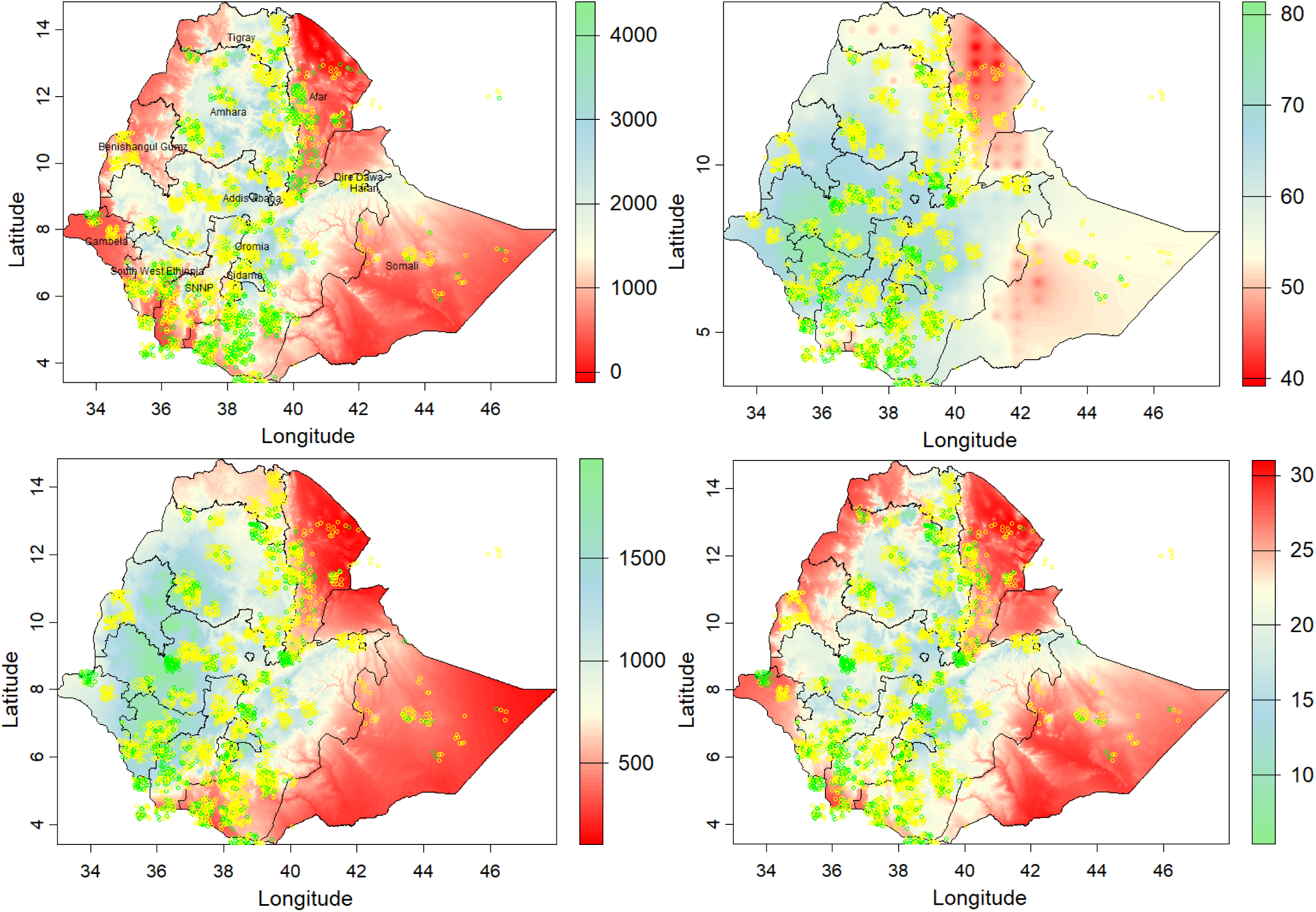
Grids of bioclimatic covariates used in the analysis of CBPP, FMD, and LSD risks in Ethiopia. Top left: altitude grid with households (data points) reporting (yellow points) and not reporting (green points) CBPP outbreak. Top right: relative humidity grid with households reporting (yellow points) and not reporting (green points) FMD outbreak. Bottom left: precipitation grid with households reporting (yellow points) and not reporting (green points) LSD outbreak. Bottom right: temperature grid with households reporting (yellow points) and not reporting (green points) LSD outbreak.

### 2.2 Model specification and fitting

#### 2.2.1 Explanatory variables

##### CBPP

To model the response variable, we reviewed the literature for environmental risk factors associated with CBPP, FMD, and LSD prevalence. A review of CBPP prevalence over the period 1996 to 2016 in Ethiopia showed that CBPP seroprevalence is highly associated with specific agroecological zones (Abdela and Yune 2017) which are determined by altitude in Ethiopia. Altitude as such is also found to be associated with the seroprevalence of CBPP (Mamo et al. 2018; Fenta et al. 2024). Mobile pastoralism and large herd sizes, which are the dominant practices in the arid and semi-arid agroecological zones in the eastern, northeastern, and southeastern parts of the country, are also identified to facilitate the transmission of CBPP. Large herd sizes as well as small herd sizes but in highly densely populated areas result in high livestock density which is a major risk factor for infectious diseases. Although interaction with or direct contact with infected animals is a primary route of CBPP transmission, aerosol transmission to a distant location (up to 200 m) can occur under favorable conditions of relative humidity and wind (WOAH, 2025). All the risk factors identified, except contact with or proximity to diseased animals and relative humidity, are highly correlated with altitude. To avoid model unidentifiability which arises when correlated explanatory covariates are included in the estimation model, we included altitude and relative humidity to explain CBPP prevalence. We included spatial effects in the model in the form the geographic coordinates at the observed locations, expecting to account for the effect of proximity to diseased herds or populations and any remaining spatial heterogeneity in disease prevalence not explained by the environmental covariates. The spatial term was included for FMD and LSD analyses as well.

##### FMD

Cattle are mostly infected with FMD by aerosol or airborne route (Zewdie et al., 2023). Risk factors in the environment and ecosystem involve factors like temperature, humidity, and the presence of virus-shedding animals. Specifically, at high RH (86%) and high temperatures (37° C), persistence (survival probability) of infectious viral particles can be expected to remain above 40% for five months on vegetation and two weeks on inanimate surfaces, suggesting that high RH provides protection for the virus when temperatures rise. We included the interaction effects of relative humidity and temperature in the estimation model.

##### LSD

Warm, humid climates, vegetation cover, and wind are favorable for biting fly populations, making them a major risk factor for LSD. The environmental covariates temperature, precipitation, and herbaceous cover were selected to explain the variability in LSD prevalence in the current study. Preliminary analysis with bioclimatic covariates (temperature and precipitation) affecting the survival and reproduction of LSD virus and vegetation cover (shrubs and herbaceous vegetation in a grid of 300^2^ meters) as covariates of LSD vector habitat suitability showed both sets of covariates equally explain the data. However, inclusion of both sets of covariates in the model resulted in insignificant effect of vegetation cover and hence removed from the final model.

Annual mean temperature and precipitation grids with a resolution of one km^2^ were collected from WorldClim (https://www.worldclim.org/data/bioclim.html). Data for relative humidity and wind speed was downloaded from NASA website (2025) as excel file, which was used to interpolate into a grid of the study area using inverse distance weighting interpolation method. Grids of land cover including herbaceous and shrubs vegetation cover were collected from Buchhorn et al. (2020). The covariate grids were then clipped by the polygon of the study area. The grid values at the coordinates of the household sampling points were extracted from the grids. The covariate grids with the households reporting or not reporting disease outbreaks are shown in Fig. 1.

#### 2.2.2 Model fitting

All analyses in this study are conducted using the R programming environment for statistical computing and analyses (R Development Core Team, 2023). We fitted generalized additive mixed models (GAMM). GAMM are a generalized linear model with a linear predictor involving a sum of smooth functions of covariates and which allow for rather flexible specification of the dependence of the response on the covariates, by specifying the model only in terms of ‘smooth functions’, rather than detailed parametric relationships (Wood, 2017). We fitted a binomial model for the binary response and ordinal model for the ordinal response datasets. The data was highly imbalanced with 76-80% of the responses being zero. Diggle and Giorgio (2015) state that prevalence data collected from community surveys often show an excess of zeros, i.e. zero-inflation, and suggest a method for count data to account for zero prevalence estimates some of which could be unreliable. For our binary data, we calculated weights for each observation to enhance reliability/accuracy. The weight was calculated as a proportion of the respondents who reported (for ‘Outbreak’ class) or did not report (for ‘No outbreak’ class) outbreak within a *kebele* (the lowest administrative sampling location, which is equivalent to a village) which is homogeneous in many aspects including climatic conditions and animal health services and where similar disease prevalence is expected in general. The weights are included in the models as a separate term.

The five surveys were conducted in different years and each survey in some of the eight regional states which are named in Fig. 1 and Fig. 7. The sampling in space was thus not identical across the five years. It is expected that the baseline disease prevalence would vary across years and locations both due to specific situations of the locations and variation in predictors, including climate variation over time. To account for the variation introduced by time-space, a factor variable combining year (2012 to 2019) and location (8 regions) was created and used as a random factor in the model. Hierarchical smooth models with random intercept varying across the fifteen levels of the year-location random factor and random smooth coefficients varying in a non-linear manner across year-location factor levels were fitted. The varying-coefficient term was modelled as the sum of a population-level global smooth relationship and year-location specific local smooth deviations from the population with ‘fs’ at smooth function using Tin Plate Regression Splines in the R package mgcv (Wood, 2025b).

The spatial smooth was modelled with two-dimensional gaussian process smooth as implemented in mgcv. The gaussian process basis function is like kriging model which is very relevant for modelling transmissible/communicable diseases as it accounts for the proximity of observations in predicting response values. The challenge with fitting models combing environmental covariates and spatial effects is the issue of spatial confounding expressed in global or local autocorrelation of residuals due to unmodelled covariates. To overcome the issue, we used NCV (neighborhood cross-validation) model estimator as suggested by Wood (2025a). First, we assessed the dependence structures in the data and determined the autocorrelation range by fitting kriging variogram models using R packages gstat (Pebesma, 2004) and automap (Hiemstra, Pebesma, Twenhöfel, & Heuvelink, 2009). Using the estimated range, we delineated for each observed location a buffer area consisting of neighboring points within the autocorrelation range using blockCV R package (Valavi et al., 2019). We then constructed a neighborhood structure using mgcv package (Wood, 2025b). Prediction of response values for each observation would be based on observations outside the neighborhood of each observation.

Below are shown the generalized additive mixed models constructed, adopting model representation from Wood (2025a). For CBPP analysis, *x_i_* is the covariate altitude, *x*_1_ and *x*_2_ represent relative humidity and wind speed as an interaction term with a tensor product thin plate spline; for FMD, *x*_1_ and *x*_2_ are relative humidity and temperature as an interaction term with a tensor product thin plate spline; for LSD *x*_1_ and *x*_2_ are temperature and precipitation as an interaction term with a tensor product thin plate spline. The *j* are smooth functions. *F* is the year-location factor variable modelled in factor-smooth interaction term with the covariates. The ƒ_0_ is the gaussian process smooth term represented by the *x* and *y* coordinates.

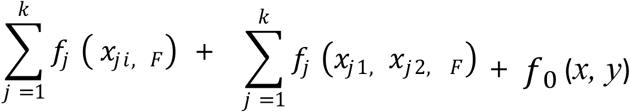

#### 2.2.3 Geostatistical mapping/interpolation

To develop disease risk maps covering the whole study area, predictions were made at the unsampled locations in the study area. Although mgcv has built-in method to predict at unobserved locations directly using the fitted GAMM model, this was not possible as data for the random year-location factor variable is not available on the predictor grids. We thus used the kriging (Cressie 1990) geostatistical spatial interpolation method using the R packages gstat (Pebesma, 2004), automap and terra (Hijmans et al., 2023). Kriging predicts at unobserved locations based on the spatial autocorrelation of observed values (Goovaerts 1997). The ability of kriging to make spatial predictions has been recognized in many studies (Aalto et al. (2013). We used a combined approach for interpolation, taking the advantages of GAMM, that it does not assume linear relationships between response and explanatory variables, and kriging, that it is a good spatial interpolator based on autocorrelation between points. We modeled variogram of fitted values from GAMM and made interpolations of the fitted values on the grid surface of the study area using kriging with external drift with the same covariates and spatial terms as used in the GAMM model. A combined or hybrid GAM-geostatistical (Aalto et al., 2013 and references therein) approaches have been used previously. Since the problem of short-range autocorrelation can be avoided, or at least mitigated, by predicting each datum when all the data in its ‘local’ neighborhood are omitted using the NCV method in mgcv (Wood, 2025b), we did not need to interpolate the residual from the GAMM model and add it to the interpolated fitted values, which is an approach recommended to address short-range autocorrelation (Aalto et al., 2013 and references therein)

The performance of the kriging interpolator model was evaluated with two measures of model fit, namely root mean squared error (RMSE) and correlation between observed and predicted values. Independence of the model residuals was assessed based on the correlation between the residuals and the predicted values which was also assessed graphically. The measures of model fit were derived from a 10-fold cross-validation. Uncertainty of the interpolation was measured by the standard error of the interpolated values.

## 3 Results

### 3.1 Model fit

Comparison of the two definitions of the response variable, i.e., as binary and four-class ordinal variable, based on the fit of the ordinal models which were fitted using REML estimation method is shown in Table 1. The model fitted for the binary response (Response2) was superior in all measures of model fit criteria (percent of the null deviance explained, AIC and RMSE) for all the three datasets of CBPP, FMD, and LSD. Subsequent results are thus presented from the analysis of the binary response data fitting the final model with NCV estimation method.

**Table 1.** Measures of model fit for two response variables defined as ordered categorical variable with four classes of disease prevalence (Response1: None, Low, Medium, and High) and two classes (Response2: disease presence or absence) fitted with REML method to estimate risks of CBPP, FMD, and LSD diseases in Ethiopia.

|  | CBPP |  | FMD |  | LSD |  |
| --- | --- | --- | --- | --- | --- | --- |
|  | Response1 | Response2 | Response1 | Response2 | Response1 | Response2 |
| Deviance explained (%) | 34.6 | 44.4 | 38.4 | 45.4 | 44.7 | 55.7 |
| RMSE | 0.746 | 0.588 | 0.661 | 0.541 | 0.720 | 0.540 |
| AIC | 1954.384 | 1251.833 | 1589.894 | 1100.434 | 1864.664 | 1099.159 |

The model fitted to the CBPP data using NCV method explained 53.8% of the model’s residual deviance. The FMD and LSD models explained 55.7% and 62.1% of the null deviance. A graphical analysis of the model residuals by plotting variogram of the residuals (Fig. 2) did not indicate the presence of unmodelled spatial autocorrelation of the residuals, since the variograms did not show sharp increase in the semivariance and plateauing after the sharp increase as would be expected for autocorrelated residuals.

**Figure 2.**
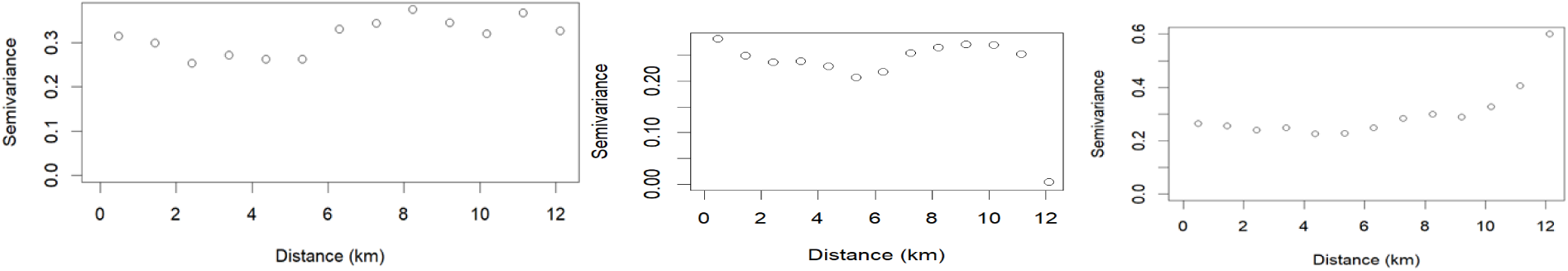
Residual variograms for assessing the dependence structures (autocorrelation) in the residuals for (from left to right) CBPP, FMD, and LSD datasets.

### 3.2 Bioclimatic and spatial effects

#### CBPP

The risk of CBPP varied significantly and smoothly with the interaction effects of relative humidity (RH) and wind speed (*P* < 0.0001; Table 2). The effects were estimated by 15 smooths corresponding to the the 15 levels of the year-location random factor (five years: 2012, 2014, 2015, 2018, and 2019 and eight regions: Afar, Amhara, Oromia, Benishangul, Gambella, Oromia, Tigray, and Somali regions). The estimates varied across the 15 factor-smooth interaction terms as smooths were modelled to allow randomly varying intercept and coefficients. The partial effects (i.e., excluding the effects of the other terms in the model) of relative humidity and wind speed are presented for selected year-location factor levels where CBPP was found most prevalent from interpolated risk maps shown in Fig. 7. The risk of CBPP is high in areas where RH is 60% or above, even under less windy conditions. The risk is very high when both RH and wind speed are high. The risk could also be high under low RH if the wind speed is high. CBPP can also be a risk in areas with low RH but high wind speed (Fig. 3). In general, the likelihood of CBPP occurrence would be up to 10 times higher under high RH and wind speed. The effects of altitude and spatial terms were not significant.

**Fig. 3.**
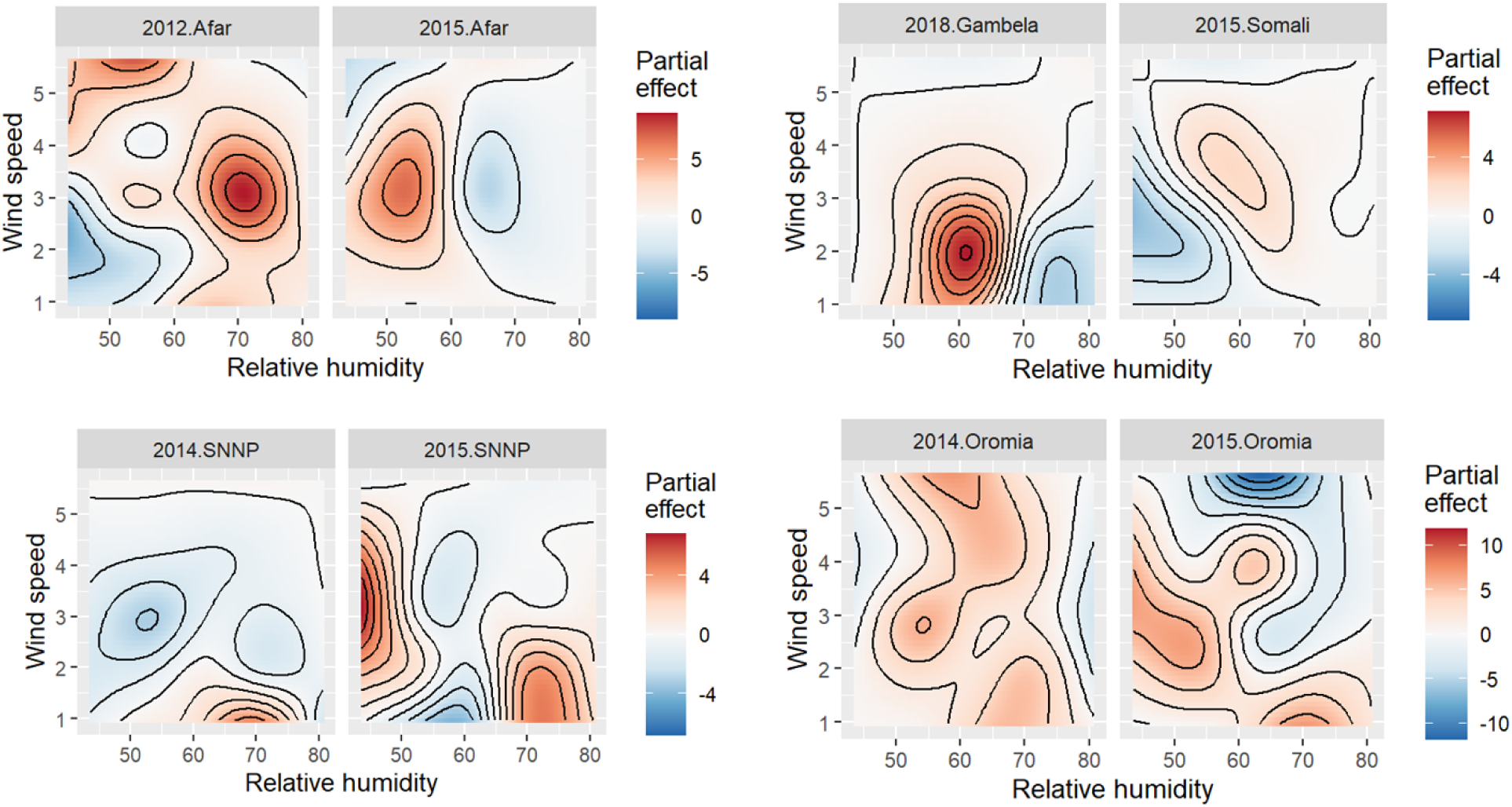
Partial effects of relative humidity (%) and wind speed (km/sec) on CBPP prevalence in Ethiopia (top left panel: smooths fitted for year.location factor levels 2012-Afar and 2015-Afar; top right panel: factor levels 2018.Gambela and 2015.Somali; bottom left panel: factor levels 2014.SNNP and 2015.SNNP; bottom right panel: 2014.Oromia and 2015.Oromia). [Prediction is over the whole range of the covariate values in the study area, and not restricted to the covariate range within the factor level]

**Table 2.** Significance of smooth terms fitted for the bioclimate and spatial effects on CBPP, FMD, and LSD disease risks in Ethiopia.

| *Smooth terms* | edf | Ref.df | Chi.sq | *P*-value |
| --- | --- | --- | --- | --- |
| ***CBPP*** |  |  |  |  |
| Altitude | 1.244 | 138 | 0.668 | 0.687929 |
| Relative Humidity*Wind speed | 68.164 | 330 | 67.238 | 0.000213 |
| Spatial effect | 120.435 | 199 | 100.515 | 0.56664 |
| ***FMD*** |  |  |  |  |
| Relative Humidity*Temperature | 45.87 | 338 | 55.62 | 0.000616 |
| Spatial effect | 99.4 | 199 | 137.41 | 8.23E-07 |
| ***LSD*** |  |  |  |  |
| Temperature*Precipitation | 60.14 | 345 | 105.03 | <2e-16 |
| Spatial effect | 82.07 | 199 | 63.04 | 0.439 |

#### FMD

The prevalence of FMD is highly determined by temperature and relative humidity (*P* < 0.001; Table 2). The risk estimated by the smooth function fitted for year-location factor level 2015.SNNP showed that the risk is very high in areas with relative humidity of ‘65-75% and temperature range of 20-25 °C (Fig.4). FMD is also highly prevalent in areas with low temperature (14-16 °C) but high relative humidity (67.5-75%) and low relative humidity (52.5-60%) but high temperature (17.5-22.5 °C) (Fig. 4; factor level 2019.Amhara). The estimate for 2019. Oromia year.location factor level also indicated high likelihood of FMD occurrence at low relative humidity (50-55%) but high temperature around 25 °C (Fig. 4). The partial spatial effects excluding the effects of the bioclimatic factors were also significant (*P* < 0.01; Table 2). The significant spatial terms indicate that the prevalence of the disease is influenced by a gaussian process as a function of distances between observations. The spatial distribution (Fig. 4) shows FMD is highly prevalent in the central and southern region of the study area.

**Fig. 4.**
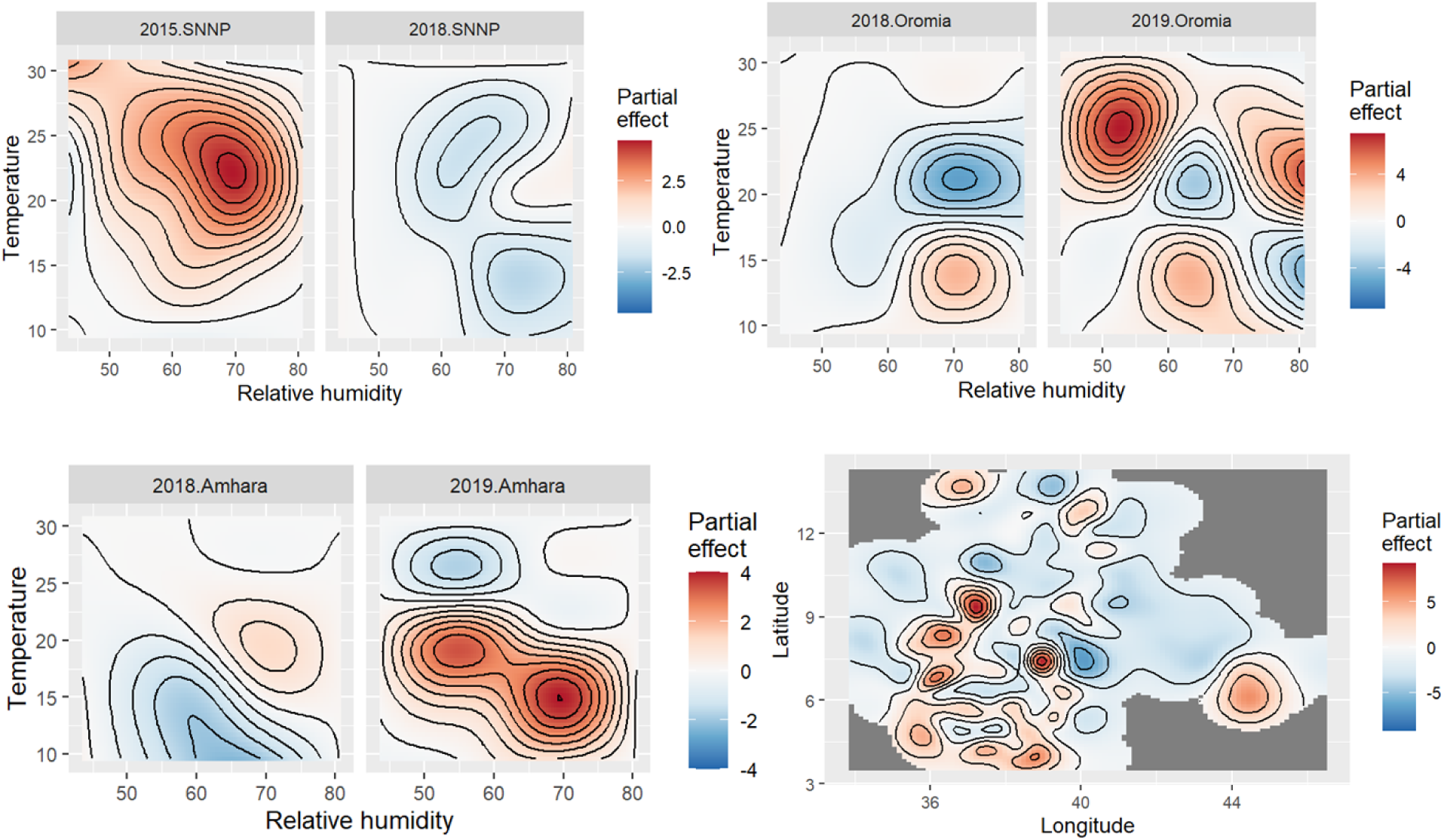
Partial effects of the interaction of relative humidity and temperature and spatial effect on FMD prevalence in Ethiopia estimated from factor smooth interaction fitted for three selected levels of the year-location (region) random factor (top left: factor levels 2015.SNNP and 2018.SNNP; top right: 2018.Oromia and 2019.Oromia; bottom left: 2018.Amhara and 2019.Amhara; bottom right: spatial effect)

#### LSD

LSD risk was found to be significantly associated with environmental covariates (Table 2). The log odds (which could be interpreted as disease risk ratio) of LSD occurrence predicted from the smooths fitted to the year-location factor level 2015.SNNP indicated that the risk could be up to 10 times more likely in areas with a temperature range of 20-30 ^0^c and precipitation of 1750-2000 mm (Fig. 5). However, the risk could also be high under low temperature (14-17 ^0^c) and high precipitation (around 1500 mm) as well as low precipitation (400-750 mm) and high temperature (22-27 ^0^c) as estimated by the smooth function for the year.location factor level 2019.Amhara. The spatial smooth plotted in Fig. 5 shows LSD is widely distributed in the study area.

**Fig. 5.**
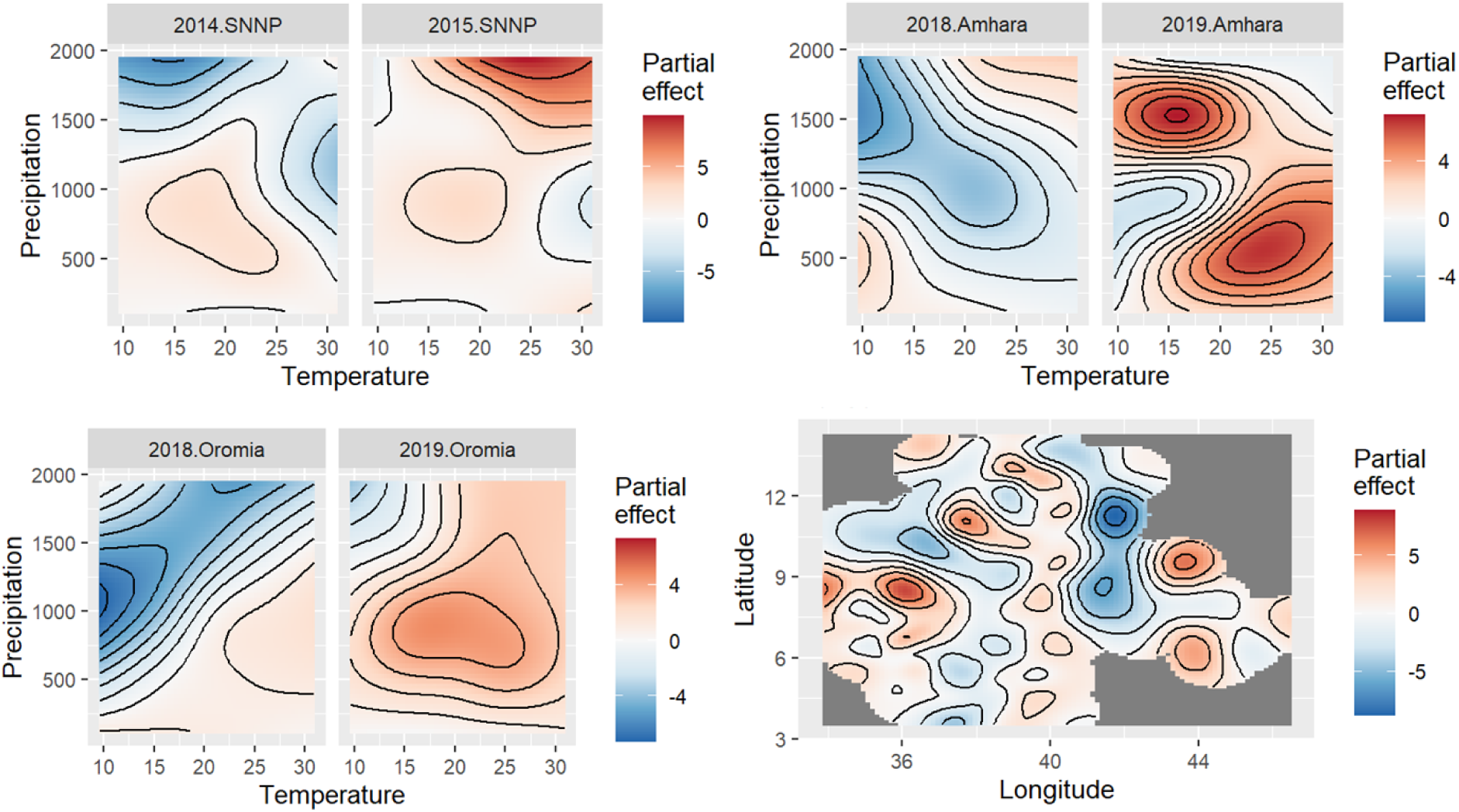
Partial effects of relative humidity (%) and wind speed (km/sec) on LSD prevalence in Ethiopia (top left panel: smooths fitted for year.location factor levels 2014.SNNP and 2015.SNNP; top right: 2018.Amhara and 2019.Amhara; bottom left: 2018.Oromia and 2019.Oromia; bottom right: spatial effects.

### 3.3 Risk maps

#### 3.3.1 Interpolator model fit

The kriging interpolator fit was found to be acceptable. The correlation between the fitted values for the observed locations from the GAM estimation model and the predicted values by the kriging model for the same observed locations is very high, being 0.9962, 0.9927, and 0.9956 for CBPP, FMD, and LSD, respectively. The RMSE are 0.2622, 0.3648, and 0.2848 for CBPP, FMD, and LSD, respectively. The model fit is also shown graphically in Fig. 6. Independence of residuals from predicted values is shown in Fig. 6 where most of the values are clustered around the origin. Normality checks for the residuals are shown on Fig. 6.

**Fig. 6.**
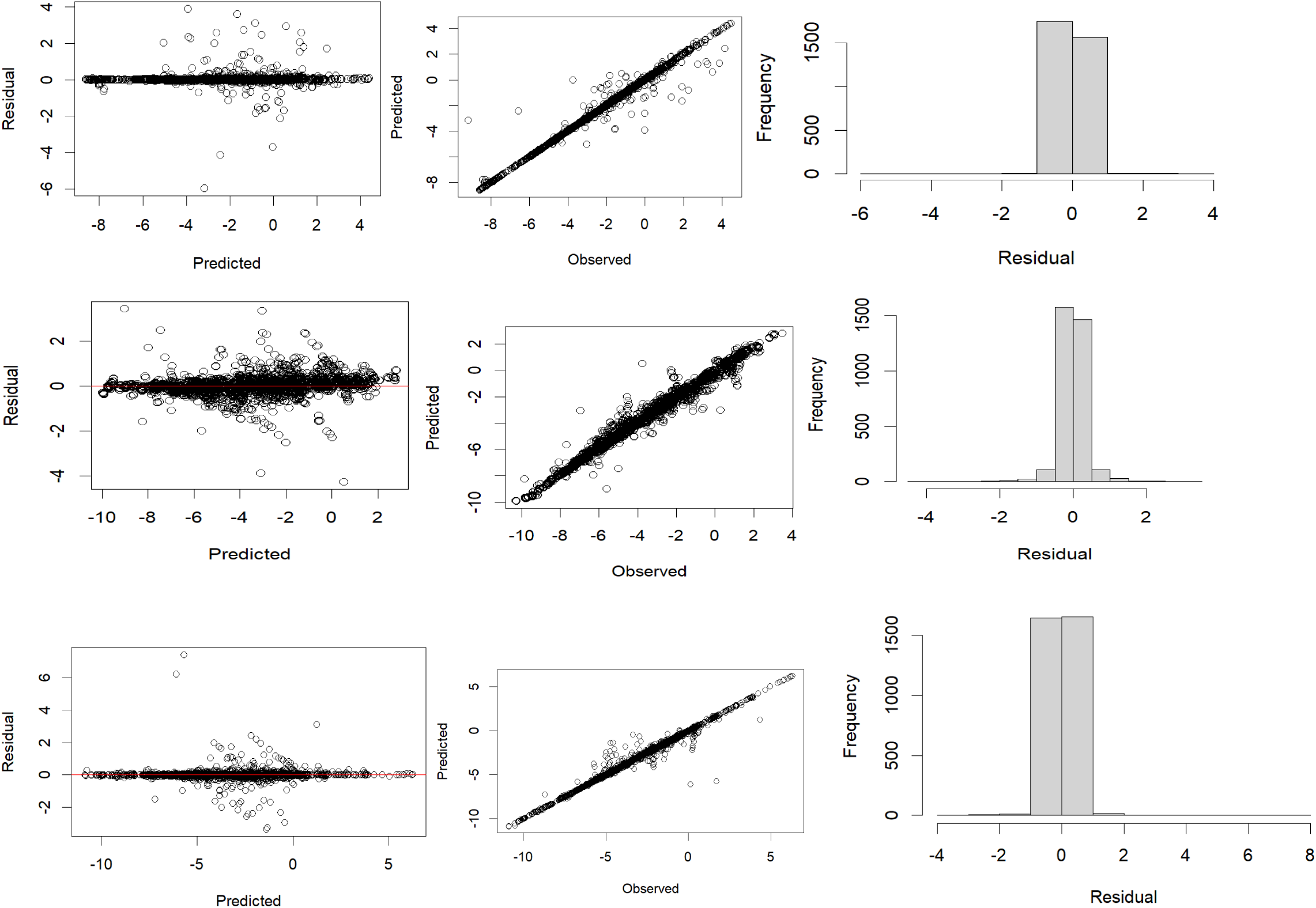
Kriging interpolator model fit (top, middle, and bottom panels for CBPP, FMD, and LSD; Left panel: independence of residuals; middle panel: correspondence between observed and predicted values, right panel: normality check for residuals

#### 3.3.2 Risk maps and uncertainties

The risk map for CBPP (Fig. 7) shows there is a higher risk in the lowland hotter areas of Ethiopia. In the areas that are colored orange and red, the odds of CBPP occurrence could be twice as high as non-occurrence. The odds of CBPP occurrence increases with increasing log odds values and declines with increasing absolute negative values, with zero corresponding to 50% probability. Fig. also indicates there could be a risk of CBPP albeit the odds is less than zero which is equivalent to less than a probability of 0.5. FMD risk is high in the central and southern regions of the study area. There is also above 50% probability of FMD risk in some areas in the north-western region. The prediction also indicated a potential risk in the western region and eastern region of Afar (Fig.8). LSD is widely distributed in the study area. The risk of LSD occurrence is up to four times likely in some hotspots shown in deep red coloring in Fig.9. Uncertainty of the predicted surface is generally low (Fig. 7 to 9). The prediction error is higher in a few border areas of the study area for all the three diseases.

**Fig. 7.**
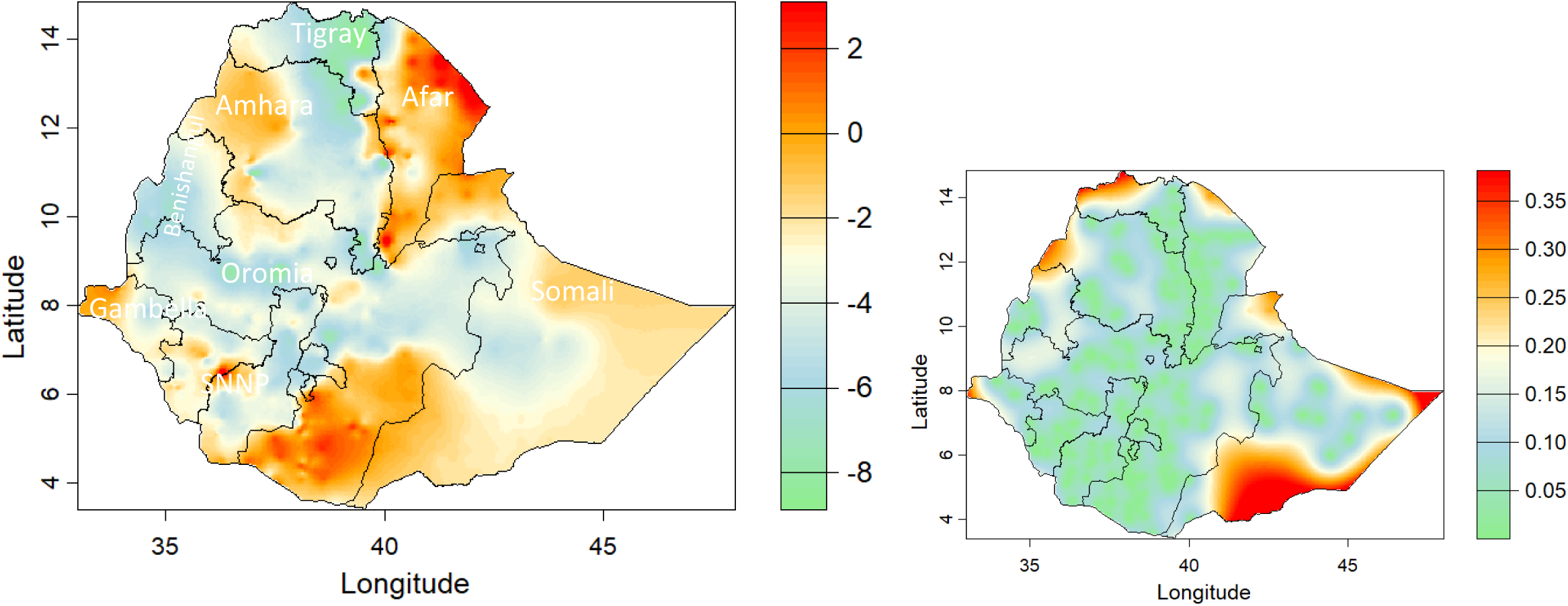
CBPP disease risk maps (right) and prediction error (right) using kriging interpolator and GAM fitted values as input data. [the risk levels are the linear predictor values in log odds scale. Positive log odds are equivalent to probability greater than 50%, negative values less than 50%, and zero equivalent to 50% probability].

**Fig. 8.**
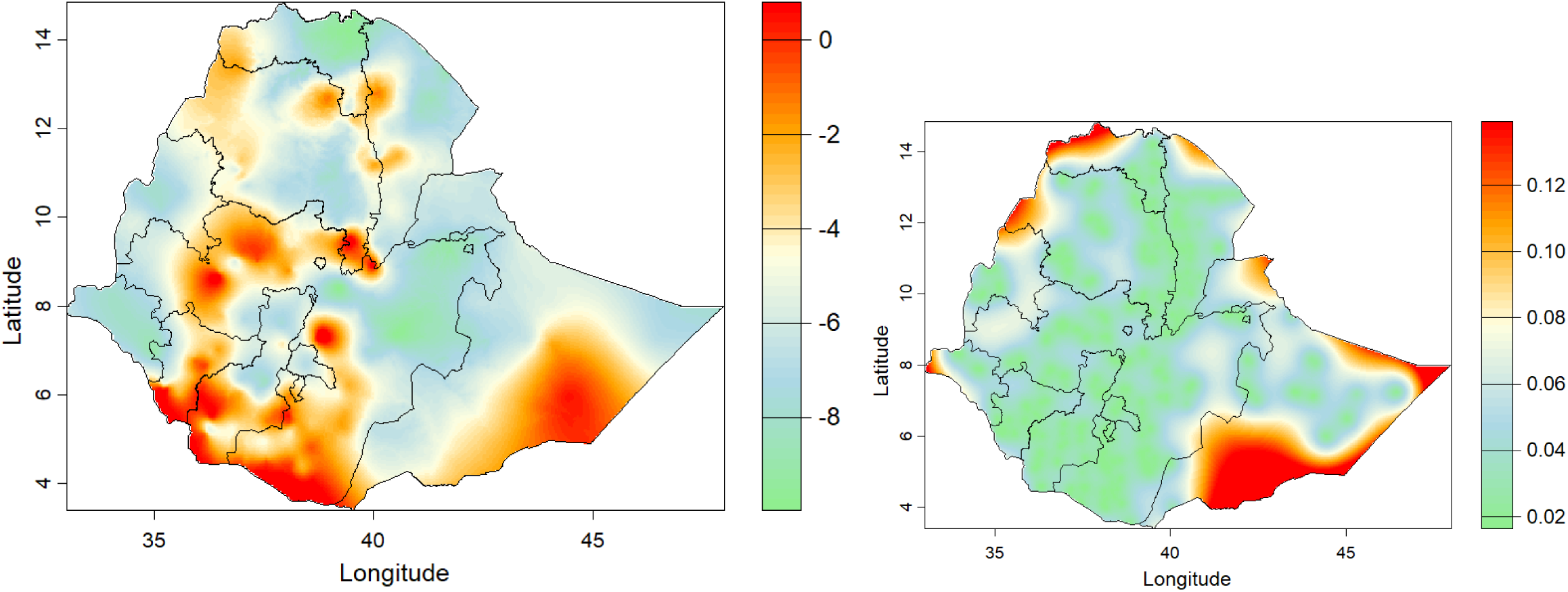
FMD disease risk maps (right) and prediction error (right) using kriging interpolator and GAM fitted values as input data. [the risk levels are the linear predictor values in log odds scale. Positive log odds are equivalent to probability greater than 50%, negative values less than 50%, and zero equivalent to 50% probability].

**Fig. 9.**
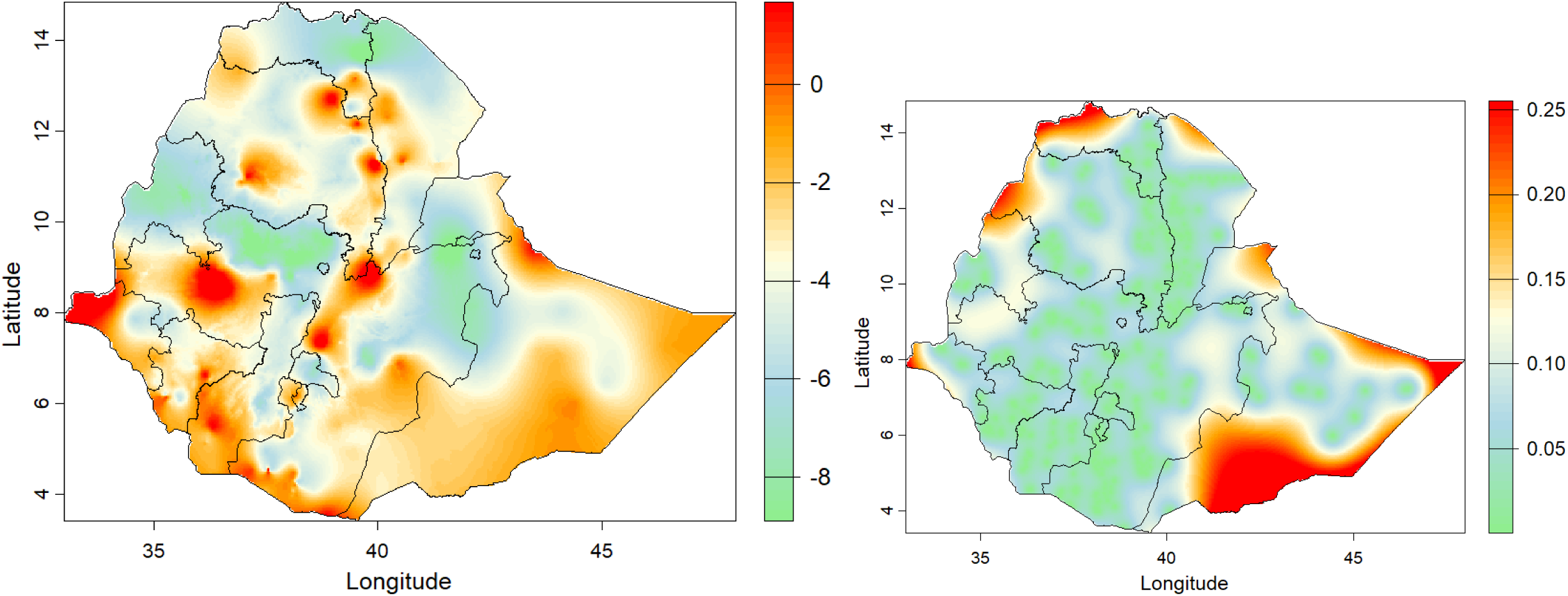
LSD disease risk maps (right) and prediction error (right) using kriging interpolator and GAM fitted values as input data. [the risk levels are the linear predictor values in log odds scale. Positive log odds are equivalent to probability greater than 50%, negative values less than 50%, and zero equivalent to 50% probability].

## 4 Discussion

### 4.1 Bioclimatic and spatial effects

In this study, using a collection of disease outbreak reports from livestock keepers’ responses to questionnaire surveys from across Ethiopia, we assessed the biogeography of three important transboundary animal diseases (CBPP, FMD, and LSD) and mapped their risks at a national scale. The models constructed with bioclimatic covariates known to affect the occurrence of the diseases fairly explained livestock keepers’ outbreak reports, as indicated by the moderate R^2^ values and the proportion of the null model deviance explained, particularly for CBPP and FMD. This indicates that livestock keepers have a good indigenous knowledge of the diseases studied here and that participatory diseases surveillance (PDS) approach, which involves community knowledge for monitoring livestock diseases using participatory appraisal methods, followed in Ethiopia is a fairly reliable alternative or complementary to the conventional epidemiology employing serological test which is infeasible to implement at a national scale due to resource limitations. The PDS approach is recognized by organizations like the World Organization for Animal Health, was of prime importance to the Global Rinderpest Eradication Program in Pakistan, was designed in the Central Asia program to provide disease information to prove freedom from rinderpest which was subsequently confirmed by laboratory serological studies (Jost et al. 2007). Questionnaire survey of experts was also used to study the epidemiology FMD outbreaks in Ethiopia (Jemberu et al., 2016). Livestock keepers in Ethiopia have vernacular names for almost all diseases affecting their livestock.

The flipside of our results, however, shows/signals that PDS results need to be integrated with geostatistical and bioclimatic modelling for a more reliable and accurate disease risk prediction. Assuming our models are accurate and that livestock populations and diseases in general are associated with bioclimatic factors, the moderate R^2^ values and the proportion of the null model deviance explained indicate PDS alone does not always suffice. While PDS is highly sensitive, allowing the detection of hard-to-find disease foci, and provides timely information, it is low in specificity and PDS results must be linked to rapid field-based test and laboratory confirmation (Jost et al., 2007). Integrating PDS results with model-based approaches could be another approach for improving the specificity of PDS results. Model-based approach also enables us to predict disease risks in unsampled/unobserved locations utilizing the association between disease occurrence and bioclimatic and other environmental variables and spatial relationships.

The smooth terms for the bioclimatic covariates assumed to explain the variation in CBPP, FMD, and LSD prevalence were all significant. Bioclimatic factors are known to be associated with diseases. Temperature, precipitation, and humidity significantly affect livestock diseases directly by causing metabolic disruptions, oxidative stress, and reduced immune function and indirectly through their impacts on quantity and quality of feedstuffs and drinking water and survival and distribution of pathogens and/or their vectors (Patel and Prajapati, 2025; Lacetera, 2019). The direct effects of climate on animal diseases are likely to be most pronounced for diseases that are vector-borne, soil associated, water or flood associated, rodent associated, or air temperature/humidity associated and sensitive to climate (Abdela and Jilo, 2016).

CBPP has been reported from the lowland areas of Ethiopia. Abdela and Yune (2017) in their review of the prevalence of CBPP over 20 years (1996–2016) found that the highest prevalence was reported from the lowland areas. Molla et al. (2021) and Kebede et al. (2022) reported a higher percentage of seropositive animals in the lowland than the midland agroclimatic zone. Mammo et al. (2018) found altitude to be significantly associated with the seroprevalence of CBPP among the potential predisposing factors assessed, with a seroprevalence of 2.5%, 3.8%, and 17.4% in the highland, midland, and lowland zones. These results agree with ours that CBPP risk increases with declining altitude. Although the pathogen does not survive for long in the environment and transmission requires close contact, under favorable atmospheric conditions of humidity and wind, aerosols can transport the agent for longer distances (WOAH website), which supports our findings that humidity does affect CBPP occurrence and transmission.

FMD occurrence and transmission is determined by the interactive effects of relative humidity and temperature (Mielke et al., 2023), with the ideal conditions for virus survival being temperatures below 50 ^◦^C, relative humidity above 55% and neutral pH (Colenutt, et al., 2020; Bartley, et al., 2002). These results partially agree with the current results where higher prevalence was estimated above 50% relative humidity for some of the factor-smooth interaction terms. For some of factor-smooth interaction levels, estimated risk was higher at higher temperature but the range of temperature in our study was much less than 50 ^◦^C. Mielke et al. (2023) also found a higher survival rate of of FMD virus at >24°C and relative humidity >75%. These authors also found that the interaction effects of relative humidity and temperature are complex with the probability of persistence of the virus in the environment decreases with increasing relative humidity at 24 °C but increases with increasing relative humidity at 28 and 31°C. Differences in the estimates of FMD risks among the 15 factor-smooth interaction levels in the current study is to be expected as the models were constructed with varying coefficients across levels. Modelling risk of airborne transmission of FMD for specific regions requires epidemiological data on circulating strains and their behaviors in aerosols (ref). Furthermore, rate of FMD virus decay varies with soil types (Bessler et al., 2024).

LSD is not well explained by our model, the R2 and deviance explained being below 50% and vegetation landcover, which is known to be a conducive environment for proliferation of the LSD vector arthropods, being not significantly associated with LSD prevalence in our study. However, the low R2 and deviance explained could also be explained by the expected low specificity of participatory epidemiology approach (Jost et al., 2007). LSD could be misidentified as respondents could have mistaken other skin diseases for LSD. LSD is significantly associated with bioclimatic and environmental variables that are suitable for the LSD vector insects and LSD virus (ref). Among the significant covariates reported by previous studies, namely precipitation, temperature, vegetation land cover, and wind speed (ref), only precipitation and temperature were found found significant factors in our study.

Climate change and extremes can directly and indirectly impact climate-sensitive diseases’ prevalence, transmission, and host susceptibility. CBPP, FMD, and LSD can be considered climate-sensitive diseases as they are significantly associated with bioclimatic variables in the current and previous studies (ref). Animal diseases that are vector borne, rodent associated, soil associated, water associated or influenced by air temperature and humidity are likely to be directly climate sensitive (Grace et al. 2015). Based on these criteria, the International Livestock Research Institute has identified a list of 38 climate-sensitive livestock diseases of most importance to vulnerable people out of 65 livestock diseases of importance to poor people (Grace et al. 2015). Temporal variation in FMD prevalence has been reported in Ethiopia (Demi et al., 2025), whereas Jemberu et al. (2016) observed neither long-term nor seasonal trends in the incidence of FMD outbreaks over a period of six years. FMD used to occur frequently in the lowland pastoral herds, but this trend has changed and currently the disease is frequently noted in the highlands of the country (Tefera, 2010 in Abdela, 2017; Seyoum and Tora, 2023). Our models constructed with random factor-smooth interaction terms with 15 levels of year-location factor also indicated varying intercepts (i.e. baseline average prevalence) and varying coefficients (i.e. varying responses to bioclimatic variables), although the factor levels do not strictly represent years because of confounding of years and locations. In the current study, we modelled bioclimatic and spatial covariates to explain the risk levels of the diseases studied. There are other animal level, socioeconomic, and production system-related factors that affect the prevalence of the diseases. These factors are well documented, for instance Jemberu et al. (2016) and need to be considered in addition to the bioclimatic and spatial indicators. One of the recommendations to address climate impacts on the livestock sector is to improve animal health service delivery (Grace et al. 2015. Access of livestock keepers to animal health services in general is low (Gizaw et al. 2021b). Efficacy of specific preventative services like vaccination are limited by the variability of and the prevalence and frequency of serotypes spatially and temporally (e.g. FMD serotypes, Jemberu et al. 2016).

The spatial terms modelled as a gaussian (kriging) process were significant for all the three diseases studied. The gaussian process modelling is a relevant approach for communicable diseases as the function explains the spill-over effect between neighboring locations within a specified distance which is determined by the autocorrelation range. The estimated autocorrelation ranges could provide valuable information for proactive emergency responses, such as imposing quarantine zones to outbreaks in neighboring regions. The kriging basis function was found superior to the other common basis function of Tin Plate Regression Spline (result not shown). This approach is relevant for diseases where cattle movement is known to be the main vehicle for transmission/spread (e.g., CBPP, Molla et al., 2017). FMD is also highly contagious and transmitted by direct contact with infected animals, with limited airborne transmission (Brown et al., 2022).The degree of spill-over effects shown by the magnitude of the effective degrees of freedoms of the spatial terms in the results section could be due to different factors – the nature of transmission of the diseases, limitation in the models fitted such as the neighborhood distance estimated, or other unobserved natural process favoring/disfavoring transmission such as seasonal movement of animals.

### 4.2 Ground truthing of Risk maps

The disease risk maps developed in this study indicate the observed geographic distribution of the diseases as reported by the livestock keepers who participated in the disease survey with potential risks predicted using a combined ecological niche and spatial modelling. Ground truthing of the risk maps was done by conducting a literature review of geographical distributions of the three diseases. All the literature reports reviewed on distribution of the three diseases are based on outbreaks reported from districts (without specific locations) most of which have different within-district ecological zones and thus different disease risk levels within district. The CBPP map from this study agrees very well with the designation of north-eastern part of the study area as the endemic zone by the Federal Epidemiology Unit in 2003 (cited in Molla et al., 2021). CBPP has been reported from the northwestern lowlands (Molla et al., 2021), southern western lowlands (Mammo et al., 2018; kebede et al., 2022), which agrees with the predicted surface in the current study. Interestingly, our risk maps show medium risk of CBPP in the central highland and northwestern lowland areas, both of which have been declared free of CBPP by the Federal Epidemiology Unit, but later refuted by (Adugna, 2017) and Molla et al. (2021).

Although FMD outbreaks are reported from all major regional states in Ethiopia, it is more frequent in the central, southern, and south-eastern parts (Jemberu et al. 2016). Based on reviews of literature published from 2008 to 2021, Zewdie et al. (2023) reported higher FMD prevalence in the southern region (Borena zone, 42.7%; Gamo zone, 26.8%) and central region around Addis Ababa (57.6% - 97.2%) than in the northern (North and South Gondar, 3.4% - 14.9%) and north-easter (Afar, 19.8%) regions. Similarly, Seyoum and Tora (2023) review of the literature from 2007 to 2020 indicate a higher FMD prevalence in Oromia region (51.8%) which spans over the central-western-southern region and Benishangul Gumuz region (19.8%) in the south-western region than in the northern regions of Amhara (7.4%) and Tigray (11.1%) and southern region (11.1%). Abdela’s (2017) concluded from his review of FMD seroprevalence until 2017 that significantly higher infection rate was observed in lowlands than in the highlands. Seyoum and Torra (2023) also found significantly higher prevalence in lowland (odds ratio = 0.67) and midland (odds ratio = 0.86) in reference to highland agroecology in their meta-analysis of literature reports from 2007 to 2020. These studies broadly agree with our predictions.

The predicted surface for LSD shows a widespread risk in study area with some hot spots in the central, southern, and north-eastern regions. Tesfaye et al. (2024) reported based on analysis of district-level LSD outbreaks in Ethiopia from 2008 to 2020 that Oromia regional state which spans across the central-western-southern region of Ethiopia reporting the highest LSD outbreaks. They also forecasted eight possible clusters of LSD outbreaks between 2020 and 2025, with five of them located in central part of Ethiopia. These results fairly agree with our risk map. It is not clear from the reports reviewed except Molla et al. (2017, 2018) whether the samplings are from herds that have not been recently vaccinated or whether the serological test results are corroborated with clinical examination of the seropositive animals. In the absence of the vaccination history of the sampled animals, it is not possible to know whether the antibodies detected are produced in response to exposure to LSD virus or recent vaccination. Some of the reports are from (semi-)intensive dairy farms which makes the ground truthing of our risk predictions which is based on survey of smallholder and pastoral livestock keepers.

In conclusion, the diseases studied here are highly associated with ecological, climatic, and spatial variabilities. These have implications for the impacts of climate change, and the vulnerability of communities in high-risk areas. Livestock disease risk mapping is an important tool for making informed policy decisions, targeted interventions, and research, but the approach is lacking in Ethiopia. The risk maps would contribute to climate-informed disease early warning systems. In the absence of model-based risk maps, disease distributions have been attributed to a whole administrative unit (districts, zones, regions) based on local outbreak reports, with under-reporting a consistent problem (Abdela and Yune, 2017; Aman et al., 2020). The maps in this study are produced using rigorous statistical analysis with very low prediction errors except relatively higher errors in the border areas and can be considered reliable.

## 6 Authors contributions

Theodore Knight – Jones: funding acquisition, project administration, writing – review & editing. Barbara Wieland: survey conceptualization and methodology, project administration, writing – review & editing. Hiwot Desta: HEARD project survey. Solomon Gizaw: HEARD project survey, data analysis, writing – original draft.

## 7 Data availability

Data will be available upon request considering ILRI’s data policy.

## 9 Acknowledgements

We acknowledge and thank the Ministry of Agriculture of Ethiopia for leading the DRSLP (I and II), RPLRP and LFSDP projects, and all the persons involved in the design of the surveys and data collection. We sincerely thank the livestock keepers who participated in the surveys. The HEARD project was financed by the European Union and implemented by the International Livestock Institute (ILRI). The DRSLP I and II projects are primarily funded by the African Development Bank (AfDB) through loans and grants, with governments contributing counterpart funds. The RPLRP and LFSDP projects are funded by the World Bank. The first, second and fourth authors acknowledge the Restoration of Livestock Systems in Drought and Conflict Affected areas of Ethiopia (RESTORE) and SAAF (Sustainable Aquatic and Animal Food Systems) projects for their research time.

